# Impact of pH and removed filtrate on *E. coli* regrowth and microbial community during storage of electro-dewatered biosolids

**DOI:** 10.1101/2021.09.07.458943

**Authors:** Tala Navab-Daneshmand, Bing Guo, Ronald Gehr, Dominic Frigon

**Author notes:** Corresponding author at: Department of Civil Engineering and Applied Mechanics, McGill University, 817 Sherbrooke Street West, Montréal, Québec, H3A 0C3, Canada.

## Abstract

Residual biosolids can be land applied if they meet microbiological requirements at the time of application. Electro-dewatering technology is shown to reduce biosolids bacterial counts to detection limits with little potential for bacterial regrowth during incubations. Here, we investigated the impacts on *Escherichia coli* regrowth and microbial communities of biosolids pH, removed nutrients via the filtrate, and inhibitory compounds produced in electro-dewatered biosolids. Findings suggest pH as the primary mechanism impacting *E. coli* regrowth in electro-dewatered biosolids. Propidium monoazide treatments were effective at removing DNA from dead cells, based on the removal of obligate anaerobes observed after anaerobic incubation. Analyses of high throughput sequenced data showed lower alpha-diversities associated with electro-dewatering treatment and incubation time. Moreover, biosolids pH and incubation period were the main factors contributing to the variations in microbial community compositions after incubation. Results highlight the role of electro-dewatered biosolids’ low pH on inhibiting the regrowth of culturable bacteria as well as reducing the microbial community variance.

## 1. Introduction

Land application of residual biosolids – produced during the secondary treatment of wastewater – has worldwide sustainability implications because of soil carbon sequestration while reducing landfill disposal demands (Lu et al., 2012). The low solids content (as low as 10-15% w/w) of raw biosolids, however, results in high transport and disposal costs (Citeau et al., 2011). Conventional mechanical dewatering processes such as belt filter press and centrifuge can increase biosolids solids content to ~30% w/w (Tuan et al. 2008), signifying the need for more effective dewatering technologies. Moreover, dewatered biosolids can be land applied if they meet certain standards. The US-EPA classifies biosolids as either Class A or B – in regards to pathogen indicators – for land application (US-EPA, 2003). Class A requirements stipulate a reduction of fecal coliforms as bacterial pathogen indicators to <1,000 most probable number per gram of total solids (MPN/g-TS), whereas Class B requires fecal coliforms <2×10^6^ MPN/g-TS. Because of this less stringent standard, land application of Class B biosolids is accompanied by a number of regulatory restrictions with respect to grazing animals and public access. An important caveat is that the above-mentioned US-EPA requirements are to be met at the time of land application and not immediately after biosolids treatment. Accordingly, Class A pathogen requirement must be achieved after transportation and storage periods, or after any mixing with other materials (US-EPA, 2003). Low bacterial pathogen counts immediately after treatment do not guarantee that the inactivation is irreversible (Dentel et al., 2008). For example, centrifuge-dewatered biosolids from mesophilic and thermophilic anaerobic digesters showed 0.8-4.1 logs increase in fecal coliforms and *Escherichia coli* after 24 h storage (Chen et al., 2011). Similarly, Higgins and colleagues reported 3.9 logs increase of *E. coli* counts to 10^8^-10^9^ MPN/g-TS after 1-2 days’ incubation in anaerobically digested centrifuged biosolids (Higgins et al., 2007). These counts then decreased back to the detection limits after 20 to 60 days.

The solids content of biosolids can be increased to as high as 70% w/w by electro-dewatering technology on a commercial scale (Zhang et al., 2017). The process uses an electrical field at the same time as pressure to remove water by electro-osmosis. Many studies have examined the parameters controlling the electro-dewatering technology’s dewatering rates, including operating factors such as constant/variable current and voltage (Qian et al., 2019), the addition of calcium oxide (Wei et al., 2020) or anthracite (Liu et al., 2021), as well as sludge properties such as apparent electrical resistivity (Sha et al., 2021a), pH, moisture content, and conductivity (Sha et al., 2021b). In addition to high dewatering rates, the electro-dewatering treatment has been shown to reduce total coliforms and *E. coli* concentrations to below the detection limits (100-600 MPN/g-TS), within 10 min, to meet Class A requirements for bacterial pathogen indicators (Navab-Daneshmand et al., 2012). Our previous work demonstrated inhibition of *E. coli* regrowth in electro-dewatered biosolids under aerobic and anerobic conditions at room temperature (Navab-Daneshmand et al., 2014a). Regrowth potentials of total coliforms and *E. coli* were compared in electro-dewatered biosolids and heat-treated controls over 7 days; no regrowth was observed in either case (Navab-Daneshmand et al., 2014a). When electro-dewatered and heat-treated biosolids were inoculated with untreated solids to mimic potential on-site contamination, counts in electro-dewatered biosolids returned to the detection limits after 4 d, whereas 4-5 logs increase were observed in heat-treated controls. The mechanisms inhibiting bacterial pathogens regrowth in electro-dewatered biosolids, however, are not clear. Regrowth of bacterial pathogens in electro-dewatered biosolids during the storage phase can be affected by several environmental conditions. The most important parameters are the availability of oxygen and other electron acceptors, temperature, water activity, pH, and the availability of nutrients. Our previous work demonstrated that the availability of oxygen as an electron acceptor has no significant impact on the regrowth of total coliforms and *E. coli* in electro-dewatered biosolids during 7-day incubations (Navab-Daneshmand et al., 2014a, 2014b). Storage temperature is another important factor that impacts bacterial regrowth, but temperature impact was outside the scope of this study and was not examined here. Water activity is a measure of water availability to microorganisms (i.e., the unbound water supporting microbial activities) and affects bacterial regrowth potentials. It is defined as the ratio of the vapor pressure of water in equilibrium with biosolids to the saturated vapor pressure of pure water. For *E. coli* regrowth to be inhibited, the water activity must be below 0.95 (Tapia et al., 2008). We have reported water activities exceeding 0.95 in electro-dewatered biosolids with total solids as high as ~60% w/w (Navab-Daneshmand et al., 2015). Therefore, water activity is not a controlling parameter for *E. coli* regrowth in electro-dewatered biosolids with the current process limitations for dewatering. The remaining two parameters and specific observations in the literature are reviewed below.

### pH

Biosolids pH affects nutrient availability and the metabolic well-being of growing bacteria. Extreme pH values cause enzymes to denature, which impairs their functions. Thus, there are pH minima, maxima, and optima for the growth of each bacterial strain. For *E. coli*, *Salmonella typhimurium*, and *Shigella* spp., the pH minima are approximately 3.90 (Koutsoumanis et al., 2004; Presser et al., 1997), and the pH optima are between 5 to 9 (Small et al., 1994). Slight changes in pH can profoundly affect regrowth dynamics; for example, *E. coli*’s growth rate increases 4.5 times from pH 4 to 5 (Presser et al., 1997). During biosolids electro-dewatering, electrolysis produces protons at the anode, and cations are electrophoretically removed with the water through the cathode, leading to a decrease in the cake’s overall pH (Navab-Daneshmand et al., 2012). It has been suggested that the lower pH of electro-dewatered biosolids (4.6 in the electro-dewatered cake compared to 7.1 in untreated biosolids) could be a factor contributing to the decrease of total coliforms regrowth (Navab-Daneshmand et al., 2014a).

### Removal of essential nutrients and precursors

The removal or addition of certain substrates or inhibitors in the biosolids matrix could impact the regrowth potential of bacterial pathogens. For example, the addition of the removed centrate from centrifuge dewatering to the dewatered cake reduced regrowth of fecal coliforms in mesophilically digested biosolids by more than 1.5 logs (Gardner and Oermeci, 2010). The removal or addition of certain autoinducers could resuscitate viable but non-culturable into a culturable state (Reissbrodt et al., 2002). In another study, the addition of substrate to mesophilically digested biosolids increased fecal coliform counts by 3.4 logs after 24 h (Chen et al., 2011). Justification for this phenomenon is that the filtrate removed during electro-dewatering could contain nutrients that are essential for bacterial regrowth, thereby reducing the regrowth of bacteria in electro-dewatered biosolids, or that the production of compounds during electro-dewatering are inhibitory to bacterial regrowth.

The synergistic effect of the above-mentioned environmental conditions could also impact the regrowth potential of bacterial pathogens. In this work we aimed to identify the impacts of pH, removed nutrients via the filtrate, and/or produced inhibitory compounds in electro-dewatered biosolids during incubation on *E. coli* regrowth and the taxonomic compositions of the microbial communities.

## 2. Materials and methods

### 2.1. Biosolids

The biosolids were obtained from an activated sludge wastewater treatment plant near Montréal. The plant receives a flow of ~60,000 m^3^/d consisting essentially of 50% municipal and 50% industrial sources (mainly pulp and paper, and food industries). It produces over 70 ton/d of residual biosolids. Two cationic polymers were added to the dewatering stream: Flo-CA475 at 1-4 kg/ton-TS before the flotation units, and Flo-CA4800 at 12-21 kg/ton-TS before the centrifuge dewatering units. Solids leaving the centrifuge units had totals solids of 17.1-19.8% w/w and pH of 6.4-7.3 with *E. coli* counts of 3.9-6.1 log MPN/g-TS. Solids samples were collected in plastic bags at the end of the solids handling unit, brought to the McGill laboratory on ice and stored at 4 °C for up to 2 d before further testing.

### 2.2. Biosolids treatment and incubation

The laboratory electro-dewatering unit used was a CINETIK® CK-lab model (Ovivo, Boucherville, Québec, Canada). The unit worked with direct current and the maximum voltage and current were set to 60 V and 5.5 A, respectively. Similarly to our previous study (Navab-Daneshmand et al., 2012), ~165 g of wet untreated biosolids were placed on the filter medium (PPS Ryton, woven) over the perforated stainless steel cathode. The ceramic-coated titanium anode applied 140 kPa constant pressure over the cake. Filtrate produced through the perforated cathode was collected in a filtrate-receiving container during the tests. The 10-min electro-dewatering cycles used in this study were sufficient to reduce *E. coli* counts to below the detection limits.

Heat treatment was used as the control treatment, because our previous results demonstrated that high temperature due to Joule heating was the main inactivation mechanism during electro-dewatering (Navab-Daneshmand et al., 2012). As previously described, 30-40 wet g of biosolids were placed in 50 mL glass tubes and incubated in a water bath at 80 °C for 16 min to achieve similar heat treatment to the electro-dewatering tests (Navab-Daneshmand et al., 2014a).

After treatment, to mimic biosolids contamination during transport or storage, some treatments were inoculated with untreated biosolids. For this, 2 wet g of untreated biosolids were dispersed in 40 mL distilled water with a homogenizer (ULTRA-TURRAX® S10N-10G, IKA Works Inc., Wilmington, NC) for 30 s. From the slurry, 1 mL was added to ~50 wet g treated sample and mixed thoroughly by hand. To control for the impact of the nutrients removed with the filtrate during electro-dewatering, in some treatments the filtrate was added back to the dewatered cake and mixed thoroughly by hand until a uniform texture was achieved. To observe the impact of pH, in some treatments HCl or NaOH solutions were added to the treated biosolids before incubation. To adjust the pH of the electro-dewatered samples to a level close to that of heat-treated biosolids, 12 mL of 0.5 M NaOH solution were added to ~50 wet g electro-dewatered biosolids and mixed thoroughly by hand. To lower the pH of heat-treated biosolids, 12 mL of 0.5 M HCl solution were added to ~50 wet g heat-treated samples. Similarly, to lower the pH of electro-dewatered biosolids with filtrate additive, 12 mL of 0.5 M HCl solution were added to ~50 wet g electro-dewatered samples that were mixed previously with filtrate.

For each treatment, approximately 50 wet g of prepared biosolids were placed in 1 L media bottles that had silicon septa on their screw caps. For uniformity, bottles were rotated horizontally on a roller apparatus during incubation (Wheaton Industries Inc., Millville, NJ). The roller operated at room temperature (22 ± 0.5 °C) and was covered to avoid light exposure. To maintain anaerobic conditions during biosolids incubation, bottles were flushed daily with pure nitrogen for 10 min. Anaerobic indicator strips (BD GasPak, Franklin Lakes, NJ) were placed in the bottles to confirm the absence of oxygen. Two sets of tests lasting 7 days each were performed 2 weeks apart.

### 2.3. Total solids, pH, and *E. coli* counts

Standard Method No 2540-B (APHA et al., 2012) was used to measure total solids. To measure biosolids pH, ~1 wet g of sample was dispersed in 9 mL distilled water with the ULTRA-TURAX® homogenizer (Navab-Daneshmand et al., 2012), and the pH was measured using a gel-filled AgCl combination electrode (Fisher Scientific, Canada).

As described in Navab-Daneshmand et al. (2012), a modified MPN method with Colilert reagent (IDEXX Laboratories, Westbrook, ME) was used to enumerate *E. coli* using Standard Method No. 9223 (APHA et al., 2012). Samples were serially diluted in clear 96-well microplates and positive well responses were measured by a SpectraMax® microplate reader (Molecular Devices, Sunnyvale, CA). An online MPN calculator was used to calculate MPN/g-TS (Curiale, 2004). The detection limit was determined to be 3 positive wells in the first row, corresponding to 60-170 MPN/g-TS.

### 2.4. PMA treatment prior to DNA extraction

Prior to DNA extraction, some of the samples were treated with propidium monoazide (PMA) to inhibit the amplification of dead or membrane compromised DNA. PMA dye penetrates cell walls and binds to the DNA bases (van Frankenhuyzen et al., 2011). When exposed to light, photolysis of the PMA dye forms a covalent bind that inhibits amplification by polymerase chain reaction (PCR) assays (Nocker and Camper, 2009). In brief, 10 mM PMA solution in 20% dimethyl sulfoxide was added to samples to achieve a final concentration of 100 μL. Samples with PMA solution were incubated in the dark for 10 min to allow sufficient binding of the PMA dye to the DNA bases. Following incubation, the samples were exposed to a 600-W halogen light at a distance of 15 cm for 5 min (Taskin et al., 2011).

### 2.5. DNA extraction and PCR

Genomic DNA was extracted from the PMA-treated and non-PMA-treated biosolids obtained on days 0 and 7 of the experiments using PowerSoil^®^ DNA isolation kits following the manufacturer’s instructions (Mo Bio Laboratories, Inc., Carlsbad, CA, USA). DNA quality and quantity were assessed using 1% agarose gel electrophoresis and spectrophotometry (260 nm/280 nm ratio). A nested PCR protocol was used to amplify 16S rRNA genes. First, DNA was amplified using universal primer pair 27F and 1492R (Weisburg et al., 1991). Next, amplicons were diluted and used as the template for a second PCR targeting the V6~V8 regions of the 16S rRNA gene using primer pairs 926F and 1392R with adaptors (Engelbrektson et al., 2010). Finally, sample-specific barcodes were added to the amplicons by using primer pairs Uniprimer1 and Uniprimer2. The primers and PCR conditions are listed in Table S1. All reactions were carried out with primer concentration at 0.3 μM, Mg^2+^ 1.5 mM, dNTP 200 μM, Taq DNA polymerase 1.25 Unit/50 μL reaction (all reagents from NewEngland Biolab). The amplicons were cleaned up using QIAquick PCR purification kit (QIAGEN), then quantified using Quant-iT™ PicoGreen™ dsDNA Assay Kit (ThermoFisher Scientific).

### 2.6. Illumina high-throughput sequencing

The amplicon library was sequenced on an Illumina MiSeq PE300 (pair-end 300 cycles) platform at McGill University and the Génome Québec Innovation Centre (Montréal, QC, Canada). In total, 3,825,375 sequences were generated as de-multiplexed forward and reverse reads. The raw sequences were paired, primer-stripped, quality-filtered, and chimera-removed using the USEARCH algorithm (Edgar, 2010), resulting in 2,220,746 good-quality sequences. Operational Taxonomic Units (I) were clustered at 97% similarity level using the open-reference protocol with the UCLUST algorithm (Edgar, 2010), assigned with 97% similarity taxonomy from the Greengenes database (McDonald et al., 2012; Werner et al., 2012), resulting in 4,026 OTUs in total.

### 2.7. Statistical analysis

Statistical analyses were performed using R statistical analysis software with the “vegan” package (R Core Team, 2019), at a significance level of 0.05. To examine the impact of different treatment factors on the regrowth of *E. coli*, an ANOVA model was applied with pH as a linear regression factor (Sokal and Rohlf, 1995). In addition to pH, biosolids treatment (heat treatment, electro-dewatering, or electro-dewatering with added filtrate), and inoculation with untreated biosolids (with or without) were also considered. The regrowth response variable was defined as the maximum log of *E. coli* counts observed during the 7-d incubation. The pH value for statistical testing was the value observed on the day of maximum *E. coli* counts; this definition was adopted because the pH increased after a few days of incubation in most of the bottles where acid was added to biosolids. Similarly, an ANOVA was applied to analyze the impact of biosolids treatment (untreated, heat treated, electro-dewatered, or electro-dewatered with added filtrate), pH, with or without inoculation with untreated biosolids, with or without PMA treatment, and before or after 7 days of incubation, on the changes in microbial community (alpha diversity). The alpha diversity indices were calculated using the number of OTUs, Hill’s number derived from the exponential of the Shannon diversity index, and Pielou’s evenness index. The effect size estimates were obtained by the omega squared (ω^2^) measure using the R “effectsize” package. The canonical correspondence analysis (CCA) was performed using the I table and five variables: pH, incubation time (days 0 and 7), inoculation with untreated biosolids, PMA treatment, and filtrate addition. The last three variables were binary. A subset of samples for CCA, including the electro-dewatered with and without added filtrate samples, was analyzed using pH, incubation time, inoculation with untreated biosolids, and filtrate addition. A heatmap of the assemblage of the top 10 genera from each sample was constructed using the R “gplots” package.

## 3. Results and discussion

### 3.1. Biosolids electro-dewatering

In this study, a typical electro-dewatering cycle started with a lag of approximately 1-min in water removal. After the onset of water removal, energy consumption and filtrate removal proceeded at constant rates (Fig. S1). Similarly, the temperature of the dewatering cake increased to ~100 °C at an almost linear rate with time. By the end of the electro-dewatering cycle (10 min), total solids increased from 18.5 ± 1.4% w/w (± indicates the range of two replicates unless otherwise specified) in untreated biosolids to 35.8 ± 0.5% w/w in electro-dewatered cakes (Fig. 1a), and the pH decreased from 6.9 ± 0.5 to 4.8 ± 0.1 (Fig. 1b). Furthermore, the 10-min electro-dewatering process reduced *E. coli* counts by at least 2.8 ± 1.1 logs MPN/g-TS, from 5.0 ± 1.1 logs MPN/g-TS in the untreated solids to the detection limit (60-170 MPN/g-TS) in the electro-dewatered cake (Fig. 1c).

**Fig. 1.**
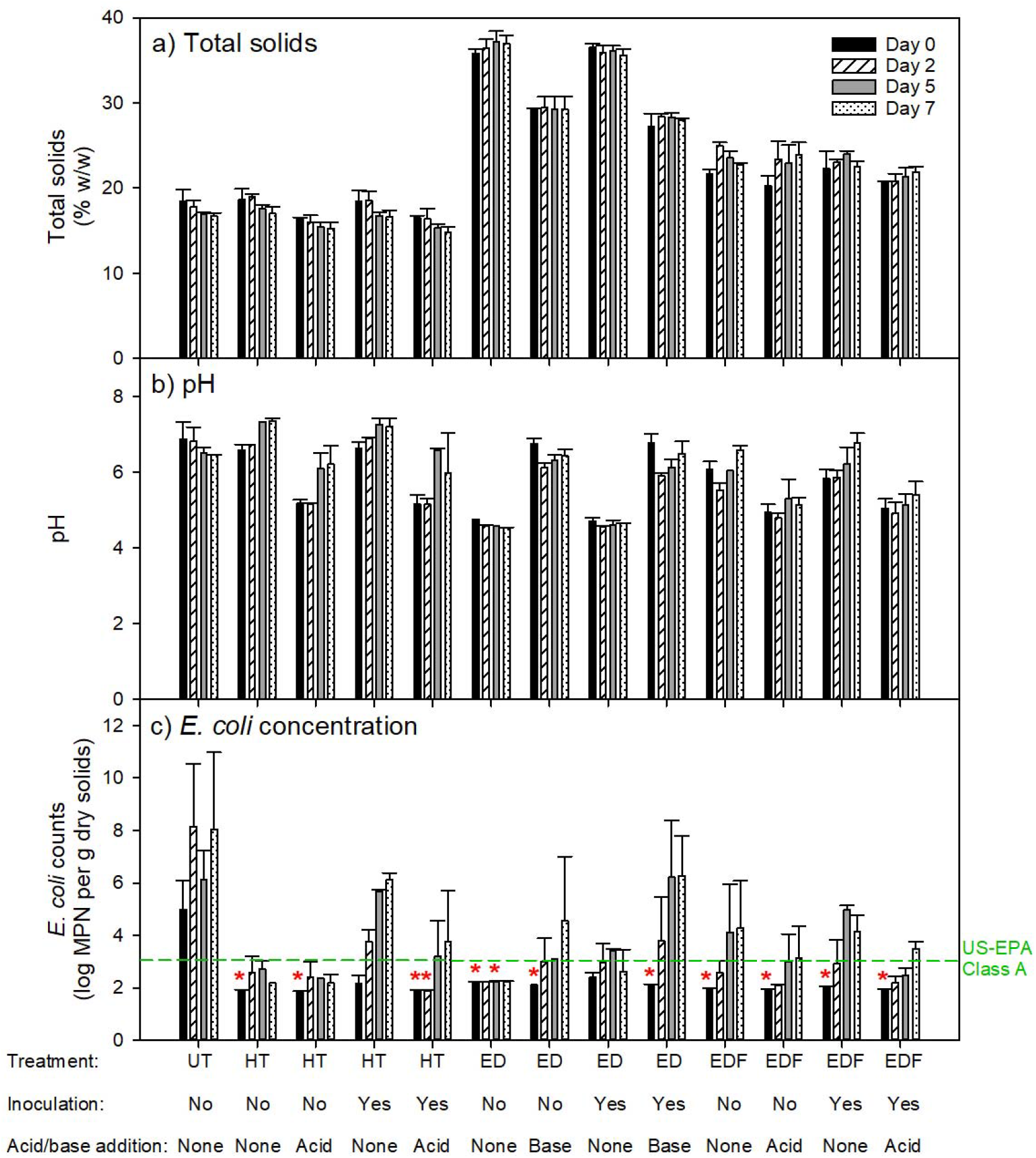
(a) Total solids, (b) pH, and (c) *E. coli* concentration in untreated (UT), heat treated (HT), electro-dewatered (ED), and electro-dewatered with filtrate addition (EDF) biosolids, with or without inoculation with untreated biosolids, and with or without acid (0.5 M HCl) or base (0.5 M NaOH) addition. Measurements were obtained during 7 d incubation under anaerobic conditions. Bars represent the range of two tests performed two weeks apart. The asterisks (*) are the detection limits (60-170 MPN/g-TS) as these counts were below detection. The US-EPA Class A threshold for *E. coli* counts is shown in green dashed line (c).

### 3.2. Total solids and pH of biosolids during incubation

The Electro-dewatering process increased the total solids content of biosolids to 35.8 ± 0.5% w/w (Fig. 1a). Addition of NaOH to electro-dewatered and inoculated-electro-dewatered samples reduced their solids content to 28.3 ± 1.0% w/w. Furthermore, addition of the removed filtrate to the electro-dewatered cake reduced total solids to 22.0 ± 0.3% w/w (Fig. 1a). The difference in total solids between untreated and electro-dewatered solids with added filtrate is due to the loss of some of the removed water by evaporation and capillarity retention on electrodes and filtration membrane. In heat-treated controls, biosolids total solids content and pH did not change from those of the untreated solids (Figs. 1a and b). In all 13 microcosms, total solids did not vary significantly during the 7- day incubation period.

Biosolids pH decreased from 6.7 ± 0.2 in untreated and heat-treated samples to 4.8 ± 0.1 in electro-dewatered biosolids (Fig. 1b). Removed filtrate had a pH of 11.8 ± 0.6, and the addition of filtrate to the electro-dewatered biosolids increased the cake pH to 6.0 ± 0.1 (Fig. 1b). During incubation, biosolids pH in samples without any acid or base additive varied over 0.2 and 1.1 pH units. In biosolids with acid or base additive, however, the pH was not as stable during the 7-day incubation period and varied over 0.5 and 1.4 pH units (Fig. 1b). There was a general increase in pH throughout the incubation period when acid had been added, while after base addition 0.6 to 0.8 pH units reduction was observed on day 2 followed by a re-increase of 0.3 to 0.6 pH units after 7 days.

### 3.3. Regrowth of *E. coli* during incubation

*E. coli* levels were reduced by 3-6 logs MPN/g-TS to the detection limits after both electro-dewatering and heat treatment (Fig. 1c). Addition of HCl, NaOH, or removed filtrate, or the inoculation of electro-dewatered or heat-treated samples with untreated biosolids, did not significantly increase the *E. coli* levels in electro-dewatered or heat-treated solids before incubation (day 0, Fig. 1c). For baseline conditions in which no filtrate, inoculum or acid/base were added, no regrowth of *E. coli* was observed during the 7-day anaerobic incubation in the electro-dewatered cake. However, when electro-dewatered samples were inoculated with untreated biosolids, *E. coli* counts increased by 1.2 ± 0.1 logs (Fig. 1c). While there was only 0.8 ± 0.4 logs regrowth in heat-treated samples, the inoculated heat-treated biosolids showed 4.0 ± 0.6 logs *E. coli* regrowth. In electro-dewatered biosolids with added filtrate, the inoculation also increased the regrowth from 2.3 ± 1.8 to 3.0 ± 0.2 logs. This is in line with our previous study where lower regrowth was observed in inoculated electro-dewatered biosolids compared to the inoculated heat-treated controls (Navab-Daneshmand et al., 2014a).

To test the impact of pH on *E. coli* regrowth, pH was manipulated for different treatments to compare treatments and evaluate different factors independently. For this, NaOH was added to electro-dewatered biosolids, and HCl was added to heat-treated and electro-dewatered biosolids with filtrate controls, with similar pH manipulations in inoculated samples. Because of the instabilities observed in biosolids pH during the incubation period, pH was used as a continuous regression variable in ANOVA analysis rather than a measured explanatory variable. Table 1 shows details of the statistical analysis results. Low pH was shown to correlate with lower *E. coli* regrowth (*p* < 0.01). Regression analysis showed an average decrease of 0.9 ± 0.4 and 1.3 ± 0.4 (± being the standard error of 12 samples) logs *E. coli* regrowth per unit reduction of pH in non-inoculated and inoculated biosolids, respectively; however, this difference was not significant. Lowering the pH with the addition of HCl in inoculated heat-treated, electro-dewatered with filtrate, and inoculated electro-dewatered with filtrate, reduced *E. coli* regrowth by 1.0 to 2.1 logs (Fig. 1c). Conversely, increasing the pH (i.e., addition of NaOH) in electro-dewatered and inoculated electro-dewatered biosolids increased regrowth by 3.3 to 3.6 logs (Fig. 1c).

**Table 1.** Results of statistical analysis on the impact of different treatment factors on *E. coli* counts in biosolids after treatment (heat-treated, electro-dewatered, or electro-dewatered with added filtrate), with or without inoculation with untreated biosolids after 7 days of incubation. Two sets of experiments (replicates) were performed two weeks apart.

| Experimental Factors | Degrees of freedom | p-value | Effect size estimate ($\omega^2$) |
| --- | --- | --- | --- |
| Treatment | 2 | 0.85 | -0.07 |
| Inoculation | 1 | 0.07 | 0.10 |
| pH | 1 | **<0.01** | 0.53 |
| Treatment * inoculation | 2 | 0.23 | 0.05 |
| Treatment * pH | 2 | 0.70 | -0.06 |
| Inoculation * pH | 1 | 0.17 | 0.04 |

The effect of biosolids treatment on *E. coli* regrowth was not statistically significant (Table 1), suggesting that neither removal of nutrients with the filtrate nor the possible production of inhibitory compounds played a major role in inhibiting regrowth. The impact of removal of essential nutrients with the filtrate can be visualized at both low and high pH levels. For high pH biosolids, inoculated electro-dewatered biosolids with NaOH addition (i.e., filtrate removed) can be compared to inoculated heat-treated controls (i.e., no filtrate removed) where similar *E. coli* regrowth occurred (6.1-6.3 logs, Fig. 1c). Likewise, for low pH solids, inoculated electro-dewatered biosolids (i.e., filtrate removed) can be compared to inoculated electro-dewatered biosolids with added filtrate and acid, and inoculated heat-treated biosolids with added acid controls (i.e., no filtrate removed); there was no significant difference between the regrowth observed in these samples (3.4-3.8 logs regrowth). Therefore, the removal of filtrate does not seem to have a meaningful impact on the regrowth of *E. coli*. The impact of inhibitory compounds that were possibly produced during electro-dewatering can be visualized by comparing inoculated electro-dewatered solids with added filtrate to inoculated heat-treated controls. Although 1-log lower *E. coli* regrowth was observed in inoculated electro-dewatered biosolids with filtrate addition (Fig. 1c), the pH was also 1 unit lower compared to inoculated heat-treated controls (Fig. 1b). Moreover, the lower regrowth in electro-dewatered biosolids with added filtrate seems to be controlled by lower pH in these solids, and therefore, could not be linked to a production of inhibitory compounds during electro-dewatering.

Results suggest that low pH of electro-dewatered biosolids is the principal parameter controlling the regrowth of *E. coli* under the discussed experimental conditions. Electro-dewatering reduces biosolids pH to 4.8 ± 0.1, which is close to the minimum pH observed for *E. coli* growth (i.e., pH = 3.9; Presser et al., 1998) (Fig. 1b). The higher *E. coli* regrowth observed in samples with higher pH values is in line with a reported increase in *E. coli* growth rates above pH 4 (Presser et al., 1997). Storage temperature of biosolids is another factor that may vary greatly over different seasons and/or between regions and can affect bacterial regrowth potentials. However, the role of storage temperature on *E. coli* regrowth in electro-dewatered biosolids was not considered in this study. Altogether, the impact of low pH on bacterial regrowth was indicated and can be applied to other biosolids treatments. Lowering biosolids pH after treatment could be beneficial for reducing regrowth of bacterial pathogens and providing optimum growth pH for certain crops and plant types. For example, crops such as berries, eggplants, and potatoes require soil pH to be between 4 and 8 (Havlin et al., 2013). It should be noted that in addition to the microbiological quality, treated biosolids should meet stringent limits of heavy metal concentrations for land application (US-EPA, 2003). When electro-dewatered biosolids are certified as Class A, the technology will be a promising method for the land application of biosolids in terms of fecal indicator bacterial content and pH levels in certain agricultural soils.

### 3.4. Changes in microbial community during incubation

Alpha diversity (within samples) of sequenced 16S rRNA gene amplicon data were calculated using the number of operational taxonomic units (OTUs, defined at 97% identity), Hill’s number, and Pielou’s Evenness index. Alpha diversity was significantly affected by the treatment of biosolids (i.e., untreated, heat treated, electro-dewatered, or electro-dewatered with added filtrate), PMA application, and incubation period (*p* < 0.05, Table 2), but not significantly affected by pH, inoculation or filtrate addition (*p* > 0.05). The interaction of biosolids treatment with pH and PMA application also showed a significant effect on the number of OTUs.

**Table 2.** Results of statistical analyses on changes in microbial community (alpha diversity) in biosolids after treatment (untreated, heat treated, electro-dewatered, or electro-dewatered with added filtrate), with or without inoculation with untreated biosolids, with or without PMA application, and before or after 7 days of incubation. The two replicates were performed two weeks apart.

| Experimental factors | Df | Number of OTUs |  | Hill's number<sup>1</sup> |  | Pielou's Evenness |  |
| --- | --- | --- | --- | --- | --- | --- | --- |
| | | *p* | $\omega^2$ | *p* | $\omega^2$ | *p* | $\omega^2$ |
| Treatment | 3 | **<0.01** | 0.64 | **<0.01** | 0.64 | **<0.01** | 0.41 |
| Inoculation | 1 | 0.27 | <0.01 | 0.64 | -0.02 | 0.69 | -0.02 |
| pH | 1 | 0.37 | <0.01 | 0.73 | -0.02 | 0.66 | -0.02 |
| PMA | 1 | **<0.01** | 0.56 | **<0.01** | 0.26 | 0.10 | 0.04 |
| Incubation | 1 | **<0.01** | 0.26 | **<0.01** | 0.21 | **0.03** | 0.10 |
| Treatment × inoculation | 2 | 0.39 | <0.01 | 0.15 | 0.05 | 0.63 | -0.03 |
| Treatment × pH | 3 | 0.05 | 0.13 | 0.06 | 0.12 | 0.12 | 0.08 |
| Treatment × PMA | 2 | **0.03** | 0.13 | 0.10 | 0.07 | 0.57 | -0.02 |
| Treatment × incubation | 2 | 0.39 | <0.01 | 0.11 | 0.06 | 0.35 | <0.01 |
| Inoculation × pH | 1 | 0.82 | -0.02 | 0.49 | -0.01 | 0.47 | -0.01 |
| pH × PMA | 1 | 0.45 | <0.01 | 0.54 | -0.01 | 0.45 | <0.01 |
| pH × incubation | 1 | 0.32 | <0.01 | 0.91 | -0.02 | 0.82 | -0.02 |
| PMA × incubation | 1 | 0.51 | -0.01 | 0.90 | -0.02 | 0.88 | -0.02 |

<sup>1</sup> Hill's number = exponential of Shannon diversity index

PMA was used to inhibit extraction and amplification of dead or compromised DNA. Comparing the microbial communities’ alpha diversity between samples with and without PMA treatment, the number of OTUs and the Hill number were reduced by 41% and 53%, respectively (Fig. 2). The distribution of the relative abundances of OTUs shows that the number of least-abundant OTUs was reduced in PMA-treated samples (Fig. S2). Previous findings showed that the majority of the diversity observed in biosolids without PMA treatment was likely due to dead cells (van Frankenhuyzen et al., 2011), suggesting that the microbial community compositions determined after PMA treatment comprise the truly regrowing species.

**Fig. 2.**
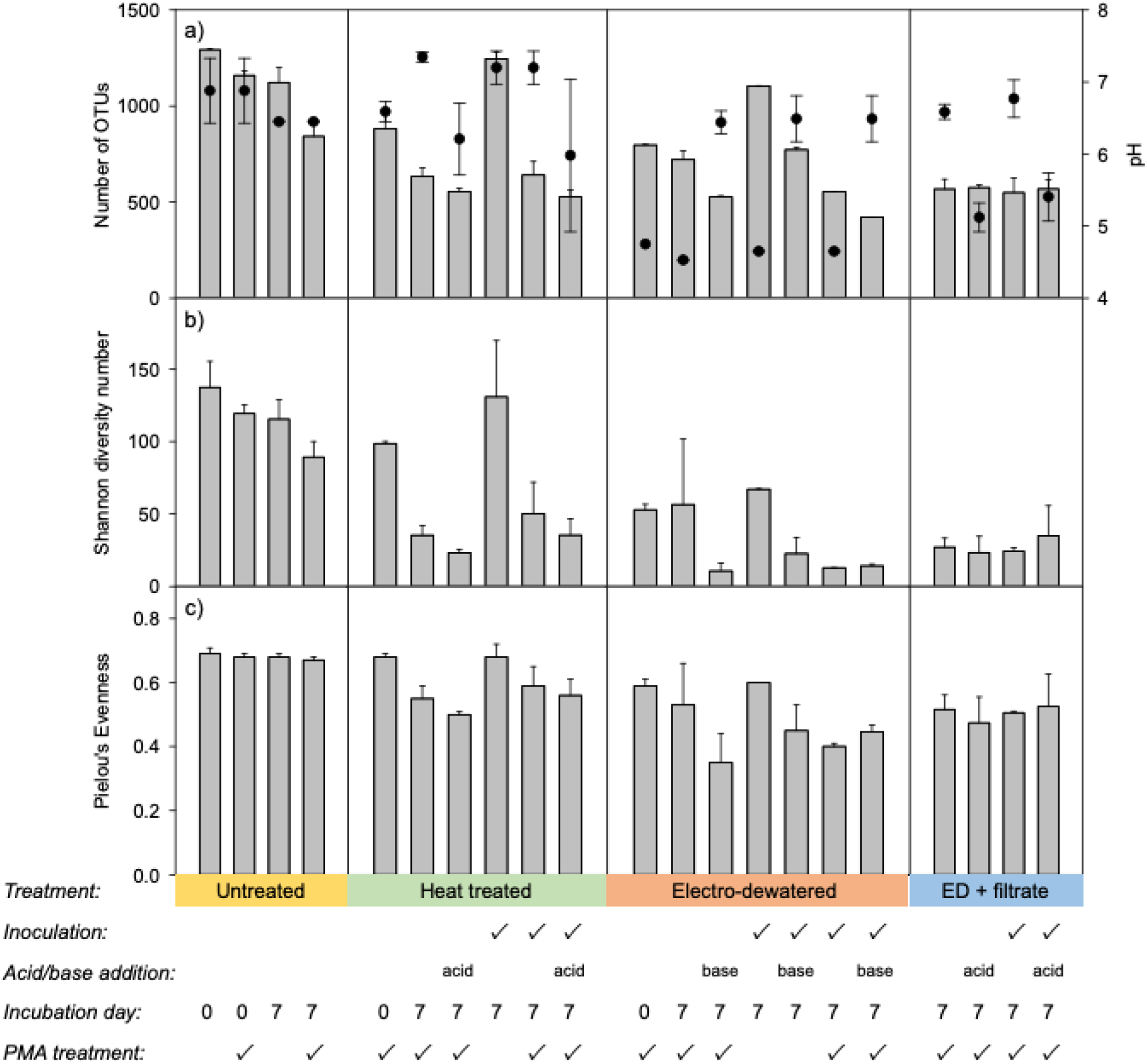
Alpha diversity indices (a) number of OTUs, (b) Hill’s number derived from Shannon diversity, and (c) Pielou’s Evenness in untreated, heat treated, electro-dewatered (ED), and electro-dewatered with added filtrate biosolids, with or without inoculation with untreated biosolids, with or without acid (0.5 M HCl) or base (0.5 M NaOH) addition, and with or without PMA treatment. Samples were collected on days 0 or after 7 d incubation under anaerobic conditions. Associated pH values (circles) are plotted on the secondary y-axis in plot (a). Bars represent the range of two replicates performed two weeks apart.

Variations in microbial community compositions with experimental factors were analyzed using Canonical correspondence analysis (CCA) to evaluate their correlations; this CCA model explained 23.4% of the variations (Fig. 3). Samples on day 7 treated with PMA tended to appear closer to the day 0 samples than the non-treated samples (Fig. 3a), but the distance between the PMA-treated and non-treated sample was relatively small. This is consistent with the observations that PMA treatment removed populations that were present at low abundance and were likely inactive. It suggests that dead cell DNA is being detected without PMA treatment. The application of PMA treatment is essential for the removal of inactive/dead cell DNA as shown in the microbial community analysis (Fig. 4). Here, approximately 50% to 75% of the least abundant OTUs were removed by PMA treatment (Fig. S2). The presence of inactive/dead cell DNA can lead to an underestimation of active/live cell DNA. For the impact of PMA application on the analysis of microbial communities, although *E. coli* species could not be identified from the 16S rRNA gene sequence data, we looked at the higher level – family *Enterobacteriaceae*. Comparing the samples with and without PMA treatment, higher abundances of family *Enterobacteriaceae* were observed in PMA-treated samples across the different pH conditions (Fig. S3). These findings show that PMA application represents a more accurate evaluation of the pH effect on the active/live microbial community. Therefore, for DNA-based detection methods (e.g., sequencing, qPCR) when both live and dead cells are present at significant levels, PMA treatment is recommended to exclude dead cell DNA.

**Fig. 3.**
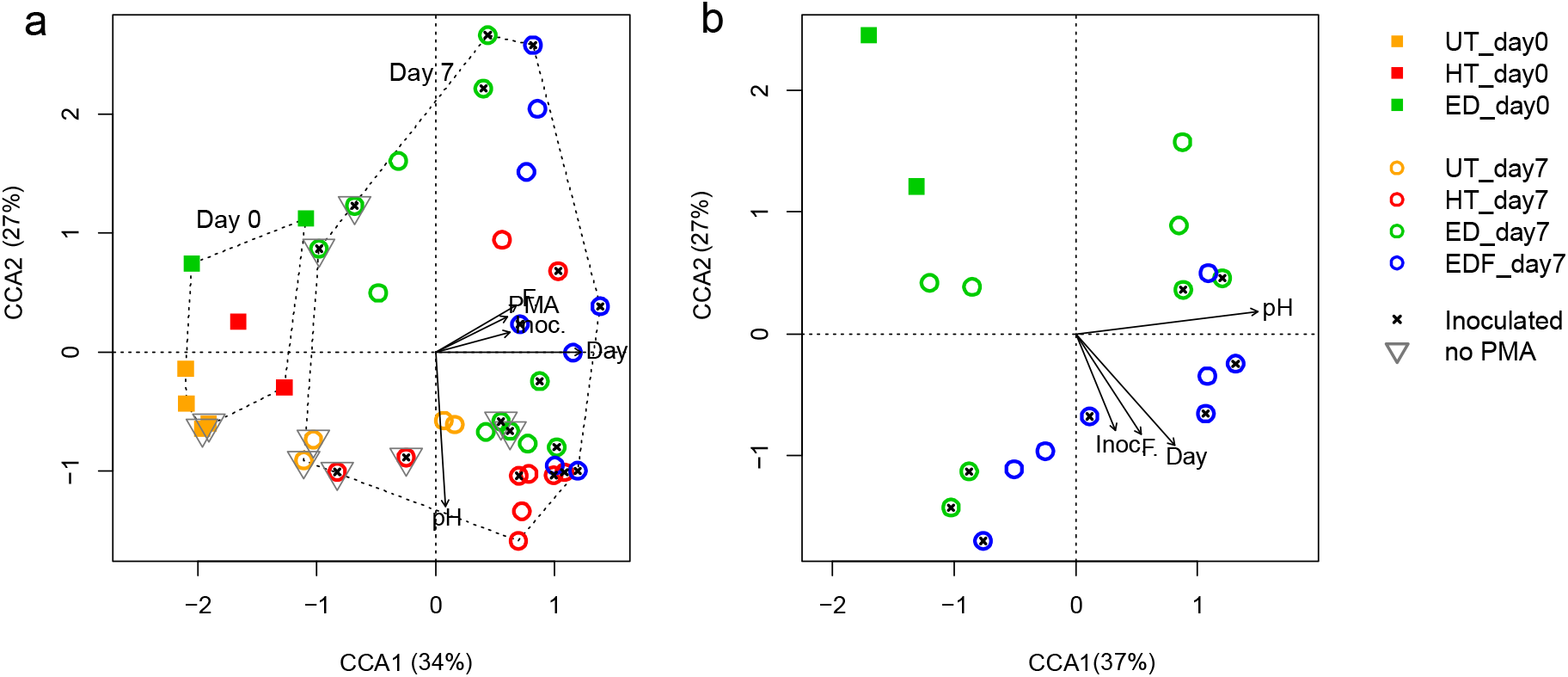
Canonical correspondence analysis (CCA) of microbial OTUs and treatment variables, pH, inoculation with untreated biosolids (Inoc.), PMA treatment, incubation time (Day), and filtrate addition (F.) in a) untreated (UT), heat-treated (HT), electro-dewatered (ED), and electro-dewatered with filtrate addition (EDF) biosolids and b) electro-dewatered with and without added filtrate. Sample replicates were from two experimental sets.

**Fig. 4.**
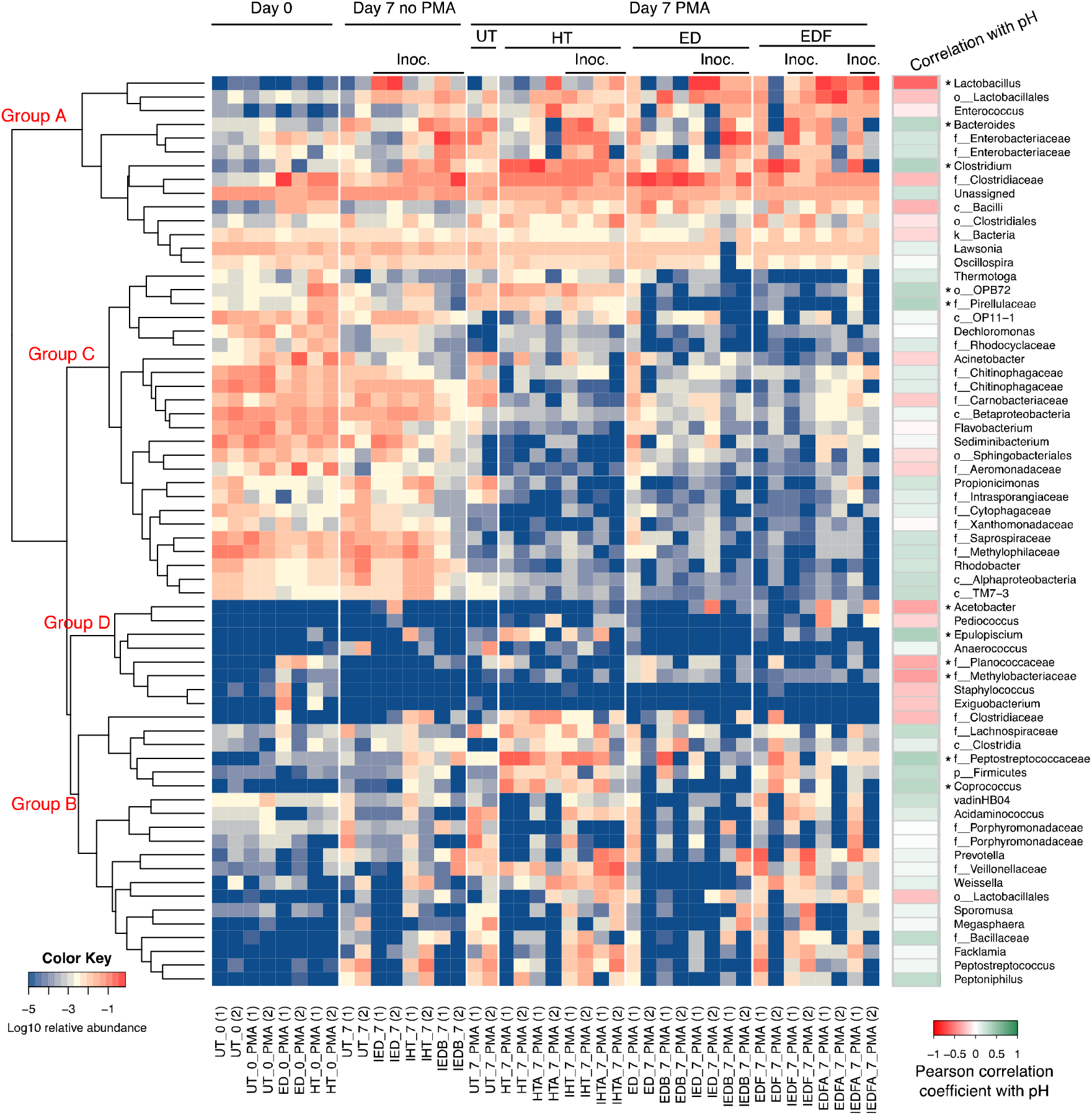
Heatmap of the top 10 genera from microbial communities and Pearson correlation coefficient with pH (* indicates *p* < 0.05). Samples are untreated (UT), heat-treated (HT), electro-dewatered (ED), and electro-dewatered with filtrate addition (EDF) biosolids, with or without PMA treatment. I, A and B stand for inoculation, acid and base, respectively. Sample replicates from two experimental sets are numbered (1) and (2). Bacteria names are shown at genus level, or higher level (family: f_; order: o_; class: c_; phylum: p_; kingdom: k_) if not identified at genus level. The colour key indicates relative abundance of genera in each sample. Hierarchical clusters indicate similarities among genera based on a Euclidean distance method.

The incubation period clearly segregated community compositions between samples on day 0 and day 7, and contributed the highest explanation power (6.6%) in the CCA model (Fig. 3a). This shows that specific OTUs grew in the community during incubation. The second factor explaining 6.1% of the microbial community variations was the final pH of the biosolids, which oriented in parallel to the CCA2 axis (Fig. 3a). Therefore, the composition of the community after 7 days of incubation was modulated by the final pH, which led to 6.1% of the variance. The remaining important experimental factors (filtrate removal, and inoculation with untreated solids) showed similar correlations to the microbial community compositions (i.e., their vector representation points in the same direction, Fig. 3a), and together accounted for 8.3% of the variations.

In order to better visualize the impact of inoculation and the addition of electro-dewatering filtrate on the community composition, a second CCA analysis was performed with only the PMA-treated ED samples. Similar to the first CCA, the incubation period and the final pH were the main explanatory factors and together they explained 20.0% of the variations. Also, the vector projection of pH was at approximately 90° from the other explanatory factors (Fig. 3b), further supporting a dominant impact of the pH on the final community profiles. The addition of ED filtrate and inoculation showed similar effects with vector projections pointed in the same direction, and they explained 7.8% and 7.0% of the variations, respectively. This observation suggests a few possible mechanisms at play. First, it is possible that the addition of ED filtrate re-introduces live bacteria, which could be similar to those introduced by inoculation with untreated biosolids. Alternatively, it is possible that the filtrate and the inoculate introduce necessary nutrients that favour certain genera.

Interpretations of the CCA analyses were refined by performing a cluster analysis on the ensemble of the 10 most abundant genera in each microbial community. This criterion selected 42 genera that accounted for between 72.3% and 98.9% of the reads in each sample. The cluster analysis identified 4 clusters of genera (Fig. 4). The last two clusters (Clusters C and D) comprised generally low abundance members of the communities in PMA treated samples on day 7. Cluster D grouped genera that were always only sporadically present in the communities. Cluster C encompassed abundant activated sludge genera and families that are either strict aerobes or respiratory facultative anaerobes with limited fermentative metabolisms (e.g., *Flavobacterium*, *Acinetobacter*, *Rhodobacter*, *Dechloromonas*, and genera in the family *Chitinophagaceae and Methylophylaceae*) (Hunter et al., 2009). These OTUs were either alive immediately after ED and heat treatments, or the cell damage was not extensive enough to allow the PMA to reach DNA. Nonetheless, they decreased in relative abundance (and likely decayed) during the 7-day anaerobic incubations for both the PMA treated and untreated samples.

Cluster A genera were the dominant populations in all biosolids samples after a 7- day anaerobic incubation irrespective of PMA treatment, but they were typically in low relative abundance on day 0, indicating their growth under anaerobic conditions. These genera included *Lactobacillus*, *Enterococcus* and other unidentified genera in the family *Enterobacteriaceae*, *Bacteroides*, and *Clostridium*. All these genera and families have highly competitive fermentative or anaerobic respiratory capabilities (Tao et al., 2020). Furthermore, they are often reported in anaerobic bioreactors (Zhang et al., 2019). Some genera in Cluster A were also abundant before the incubation (e.g., *Lawsonia* and *Oscillospira*), but these are also anaerobic bacteria found in human or animal intestines (Gophna et al., 2017). They are potentially carried by wastewater and remain in the activated sludge (Cai et al., 2014; Guo et al., 2019), indicating their ability to survive under various conditions.

Cluster B genera were generally at low abundances on day 0, and increased in abundance by day 7 of anaerobic incubation except in the sample with only ED treatment (i.e., not ED plus filtrate addition). Similar to Cluster A, the genera in Cluster B have extensive anaerobic respiration and highly competitive fermentative capabilities (Tao et al., 2020). In fact, several OTUs in Cluster B are closely related phylogenetically to OTUs in cluster A. The reduced ability of these OTUs to grow in ED treated biosolids suggest that electro-dewatering removed essential nutrients for the growth of these organisms or that inhibitory compounds that are neutralized by the return of the ED filtrate are produced.

Since pH was shown to largely affect the microbial community variance, the most abundant genera were tested for correlation between their relative abundances and pH values (Fig. 4). A few genera have been significantly negatively correlated with pH (Pearson correlation coefficient, *p* < 0.05). *Lactobacillus* and *Acetobacter* were reported in anaerobic digestion systems and related to production of volatile fatty acids (Xu et al., 2014), suggesting their favoured growth under low-pH conditions. Many species of *Lactobacillus* may produce antimicrobial substances (e.g., organic acids, bacteriocins) that can inhibit *E. coli* growth (Caridi, 2002; Drago et al., 1997). It is possible that the low-pH conditions created a suitable growth niche for *Lactobacillus* which then generated chemicals to strengthen the inhibition of *E. coli* regrowth. However, this mechanism needs to be further investigated under the biosolids regrowth conditions, which may be used as a strategy for controlling *E. coli* regrowth.

*Bacteroides*, *Clostridium*, o_OPB72, f_*Pirellulaceae*, *Epulopiscium*, f_*Peptostreptococcaceae*, *Coprococcus* have shown significant positive correlation with pH, most of which were reported in basic conditions. *Bacteroides* and *Clostridium* are core members found in many anaerobic digestion systems (Tao et al., 2020) which are typically maintained at neutral or slightly basic pH conditions. O_OPB72 was isolated from Yellowstone hot spring sediments with a pH of 7.6 at 20 °C (Hugenholtz et al., 1998). Some *Pirellulaceae* strains were isolated from a natural environment at pH > 7 (Dedysh et al., 2020; Kumar et al., 2021). *Peptostreptococcaceae* are fermentative anaerobes with optimum pH 7.5-8.3, except for the genus *Tepidibacter* (6.0-7.3) (Rosenberg et al., 2014). These results indicate that pH can affect the relative abundance of dominant OTUs due to their different optimum pH conditions for growth, while these OTUs may have little adverse effect on the regrowth of *E. coli*.

## 4. Conclusions

Mechanisms of *E. coli* regrowth and changes in microbial communities in electro-dewatered biosolids were investigated in a microcosm study. The impacts of pH, removed filtrate, and/or produced inhibitory compounds on *E. coli* regrowth and microbial communities were assessed. Results indicate pH as the primary parameter that controls *E. coli* regrowth in electro-dewatered biosolids. Analyses of high throughput sequencing showed lower alpha-diversities associated with electro-dewatering treatment and incubation time. Biosolids pH and incubation period were the main factors contributing to variations in microbial community compositions. The findings highlight the role of biosolids pH on both regrowing culturable bacteria and overall microbial communities.

## Acknowledgements

This research was funded by the Natural Sciences and Engineering Research Council of Canada’s Collaborative Research and Development program and Ovivo. We thank Frederic Biton (Ovivo), Bruno Desmarais (Ovivo), Alain Silverwood (Ovivo), Ceĺine Gagnon (Aquatech), and Gilbert Samson (Reǵie d’Assainissement desEaux du Bassin LaPrairie) for their technical support and useful comments.

