## Supplementary information for "Impact of pH and removed filtrate on *E. coli* regrowth and microbial community during storage of electro-dewatered biosolids"

817 Sherbrooke St. West, Montreal, Quebec H3A 0C3, Canada

^c^ Centre for Environmental Health and Engineering, Department of Civil and Environmental Engineering, University of Surrey, Guildford, Surrey GU2 7XH United Kingdom

^*^ Corresponding author at: Department of Civil Engineering and Applied Mechanics, McGill University, 817 Sherbrooke Street West, Montréal, Québec, H3A 0C3, Canada. Email address:

Supplementary information includes:

1 Table

2 Figures

Table S1. Nested PCR primers targeting 16S-rRNA and conditions

| Primer | Primer sequence (5’-3’) | Conditions | Cycle No. |
| --- | --- | --- | --- |
| First PCR |  | Initial denaturation: 94 ºC, 5m  Denaturation: 94 ºC, 30s  Annealing: 55 ºC, 60s  Elongation: 72 ºC, 90s  Final elongation: 72 ºC, 10m | 30 |
| 27F | AGAGTTTGATCMTGGCTCAG |  |  |
| 1492R | GGTTACCTTGTTACGACTT |  |  |
| Illumina PCR |  | Initial denaturation: 94 ºC, 3m  Denaturation: 94 ºC, 30s  Annealing: 62 ºC, 45s  Elongation: 72 ºC, 1m  Final elongation: 72 ºC, 10m | 15 |
| 926F | CTTTCCCTACACGACGCTCTTCCGATCTAAACTYAAAKGAATTGRCGG |  |  |
| 1392R | GTGACTGGAGTTCAGACGTGTGCTCTTCCGATCTACGGGCGGTGTGTRC |  |  |
| Barcode PCR |  | Initial denaturation: 94 ºC, 3m  Denaturation: 94 ºC, 30s  Annealing: 59 ºC, 20s  Elongation: 72 ºC, 45s  Final elongation: 72 ºC, 5m | 15 |
| Uniprimer1 | CAAGCAGAAGACGGCATACGAGATCGATGTGTGACTGGAGTTC |  |  |
| Uniprimer2 | CAAGCAGAAGACGGCATACGAGAT-index-GTGACTGGAGTTC |  |  |

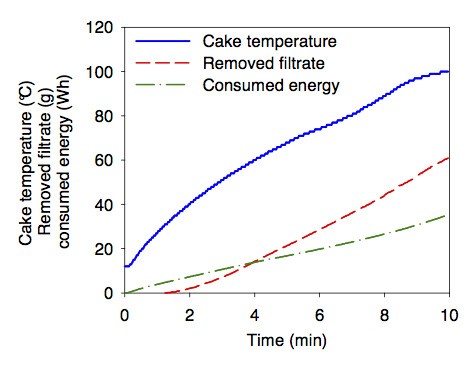

Figure S1. Cake temperature, removed filtrate, and consumed energy during biosolids electro-dewatering. The plots are averages of eight tests, performed two weeks apart with standard errors below 2.0, 4.0, and 1.7% after 4 min for cake temperature, removed filtrate and consumed energy, respectively.

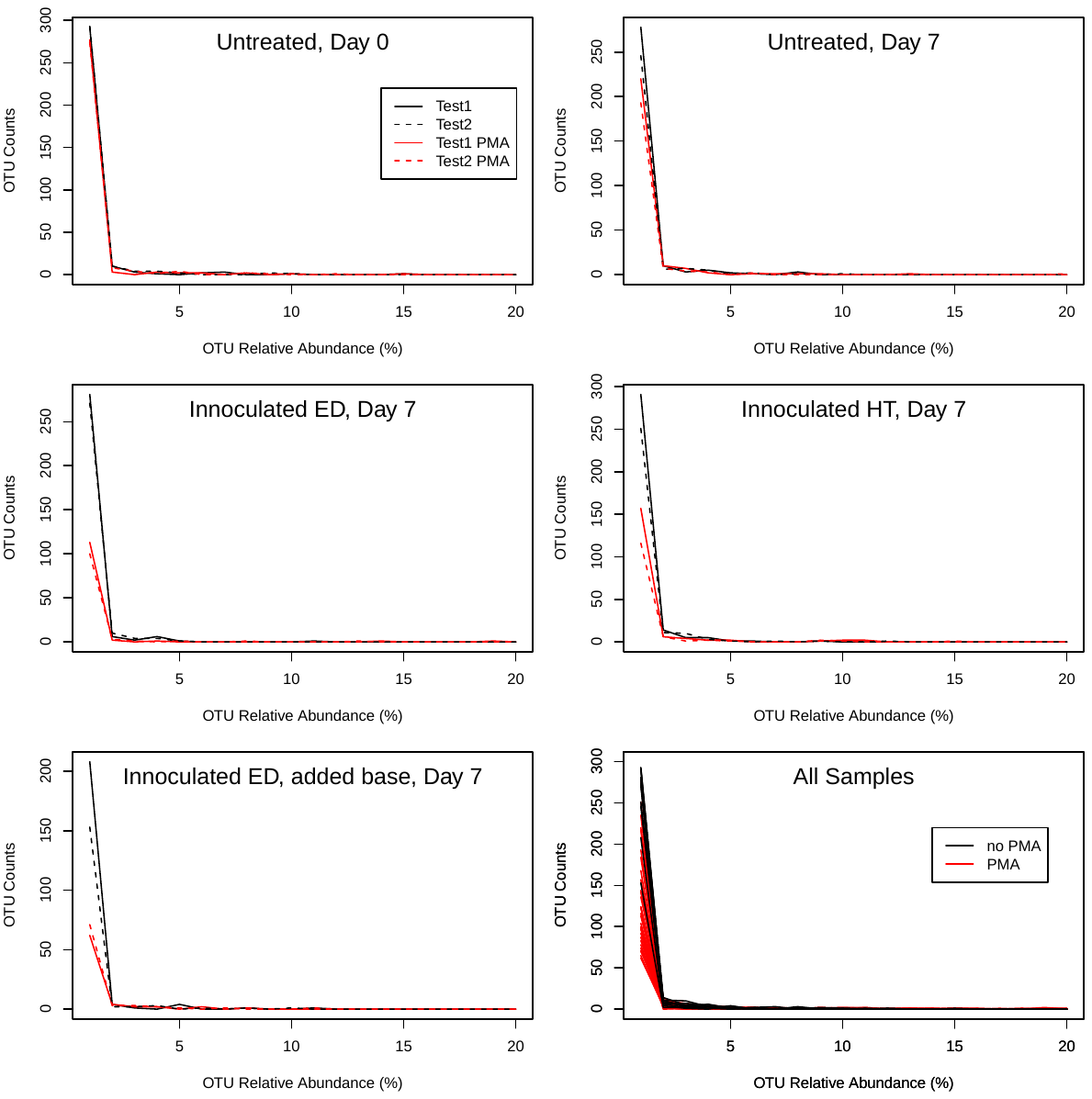

Figure S2. The number of OTUs at different relative abundances (%). Comparison between samples with and without PMA treatment in untreated, inoculated electro-dewatered (ED), and inoculated heat-treated (HT) biosolids is shown with two replicate experiment tests.

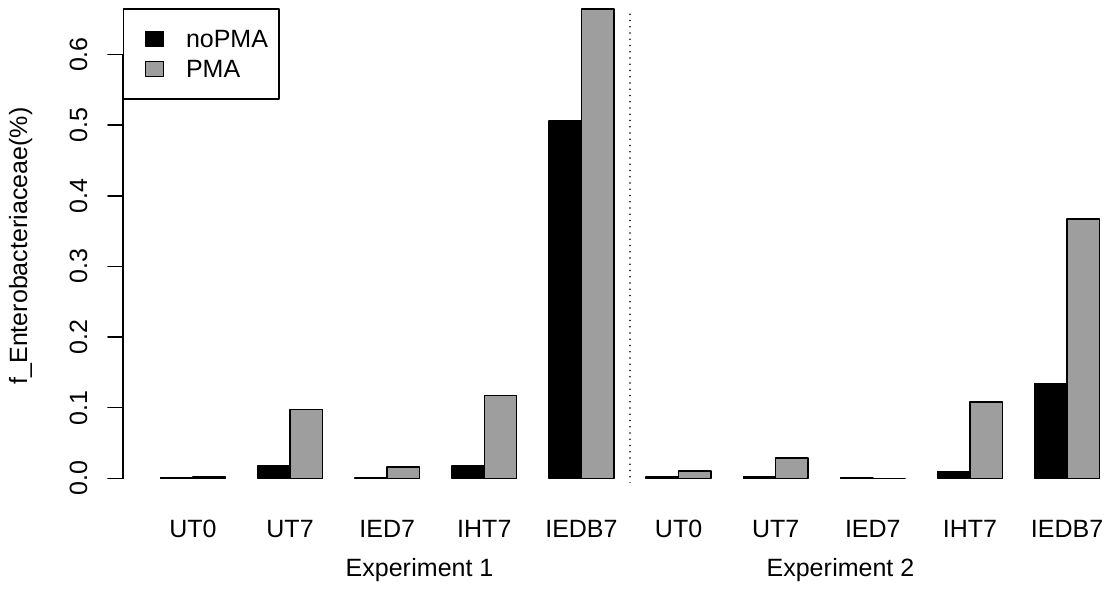

Figure S3. Relative abundance of the family *Enterobacteriaceae* with and without PMA treatment in untreated (UT), inoculated electro-dewatered (IED), inoculated heat-treated (IHT), and inoculated electro-dewatered with added base (IEDB) on days 0 or after 7 d incubation under anaerobic conditions. Data are shown for the two replicated experiments.
